# Scaling laws of graphs of 3D protein structures

**DOI:** 10.1101/2020.08.11.246041

**Authors:** Jure Pražnikar

## Abstract

The application of graph theory in structural biology offers an alternative means of studying 3D models of large macromolecules, such as proteins. However, basic structural parameters still play an important role in the description of macromolecules. For example, the radius of gyration, which scales with exponent ~0.4, provides quantitative information about the compactness of the protein structure. In this study, we combine two proven methods, the graph-theoretical and the fundamental scaling laws, to study 3D protein models.

This study shows that the mean node degree of the protein graphs, which scales with exponent 0.038, is scale-invariant. In addition, proteins that differ in size have a highly similar node degree distribution, which peaks at node degree 7, and additionally conforms to the same statistical properties at any scale. Linear regression analysis showed that the graph parameters (radius, diameter and mean eccentricity) can explain up to 90% of the total radius of gyration variance. Thus, the graph parameters of radius, diameter and mean eccentricity scale with the same exponent as the radius of gyration. The main advantage of graph eccentricity compared to the radius of gyration is that it can be used to analyse the distribution of the central and peripheral amino acids/nodes of the macromolecular structure. The central nodes are hydrophobic amino acids (Val, Leu, Ile, Phe), which tend to be buried, while the peripheral nodes are more hydrophilic residues (Asp, Glu, Lys). Furthermore, it has been shown that the number of central and peripheral nodes is more related to the fold of the protein than to the protein length.

## Introduction

Proteins, molecules that serve many critical roles in nature, consist of complex systems of amino acids that have increasingly been modelled as networks over the last decade (1–3). There are many ways to abstract the 3D protein structure into a graph. We can consider Cα, Cβ or all heavy atoms to construct an adjacency matrix. A typical method to abstract the protein model into a graph is to consider Cα atoms with 7.0 Å cut-off distance (4). We should be aware that some information is lost when the 3D model is abstracted into the graph, although it still captures relevant biochemical properties of the “real” 3D protein model. Once the 3D model of the protein is abstracted into a graph, we have several options to analyse the 3D structure by examining different parameters of the graph. Residue network models have been used to predict catalytic sites (5–8), to study protein dynamics (9), to discover node-amino acids that play crucial roles in protein folding (10,11), to explore allosteric pathways (12–14), and to analyse enzyme domain packing (15). In addition, graph theory has also been successfully used to validate PDB entries (16), to study local errors in protein models, and to discriminate decoys from the native structure (17,18). We should emphasize the importance of validation and quality assessment, since only correct protein structures can answer relevant biological questions (19–22).

However, we cannot expect that all graph parameters used and derived by mathematicians will have practical implications, for example, when studying specific phenomena in structural biology. However, we should try to find a connection between theoretical graph parameters and biochemical phenomena that can lead to deeper insights.

Despite the obvious usefulness of graph theory for the analysis of protein structures, basic structural parameters still serve an important role in the description of macromolecules. For example, a common parameter used to describe the compactness of protein models is the radius of gyration (23–25). The radius of gyration of a macromolecule describes the distribution of atoms around the centre of mass. Since Flory’s theory (26), the scaling law between the radius of gyration and the length of the protein has been studied in detail and used to describe protein folding, and to analyse the compactness of protein structures in poor or good solvent conditions. It was found that the radius of gyration of globular protein structures, including monomers and oligomers, scales with exponent ~0.4 (27). The radius of gyration can be used as a constraint when building protein models or performing molecular dynamics simulations. On the other hand, when the 3D model of a protein is not known, the scaling law provides qualitative information about the dimensions of the macromolecule.

This study links two well-established approaches, the graph-theoretical and the fundamental scaling laws, to study 3D protein models. In this paper, the PDB is surveyed to study scaling laws of 3D protein structures using a graph-theoretical approach. This research has demonstrated that the mean node degree of the protein graph is nearly scale-invariant. Additionally, the comparison of the node degree distributions of proteins of different sizes exhibits marginal differences. Furthermore, this study shows that the compactness of the protein, which is conventionally calculated by the radius of gyration, can be estimated using graph eccentricity, which also provides insights into central (buried) and peripheral (non-buried) amino acids.

## Methods

### Dataset

The Protein Sequence Culling Server (28) was used to obtain the PDB id list for protein structures with the following characteristics: maximum mutual sequence identity of 80%, X-ray resolution cut-off of 3.0 Å and minimum (maximum) chain length of 40 (10,000) residues. The PDB id list was then used to retrieve 31,571 Biological Assemblies from the Protein Data Bank.

### Graph Construction

From each of the 31,571 3D Biological Assemblies, the graph was constructed and analysed. Cα atoms were considered as nodes, and edges between nodes were constructed if the Cα–Cα distance between a pair of residues was less than (or equal to) 7 Å. It follows that the number of nodes was equal to the number of residues (Cα atoms) in the protein. Ligands, water molecules, and other hetero-compounds were discarded during graph construction. Thus, if a protein has *n* residues, then a protein graph G = G(V, E) consists of a set of vertices (nodes) V = v_1_ v_2_, … v_n_ and a set of edges E = e_1_, e_2_, … e_m_.

### Graph parameters

The mean node degree (MND) of a graph G is expressed with the ratio

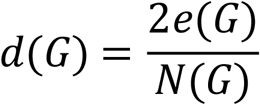

where e(G) represents the total number of edges in a graph G, and N(G) is the number of nodes in a graph G.

The eccentricity is a node centrality index defined as the maximum distance between a vertex to all other vertices. Thus, the vertex’s eccentricity is the maximal shortest path between the vertex and all other vertices. Mean eccentricity is expressed as an average value of eccentricities of all vertices of G. The radius of the graph is defined as a minimum eccentricity among all vertices in the graph. Meanwhile, the diameter is defined as maximum eccentricity among all vertices in the graph. The center of a graph or central node has eccentricity equal to the radius. A vertex is said to be a peripheral node if its eccentricity is equal to the diameter.

Next, R script (igraph package) was used to calculate mean node degree, eccentricity, mean eccentricity, radius and diameter:

~~~
MND <- mean(degree(G)) # mean node degree of graph G
EC <- eccentricity(G) # eccentricity of graph G
MEC <- mean(EC) # mean eccentricity of graph G
R <- min(EC) # radius of graph G, or min eccentricity of graph G
D <- max(EC) # diameter of graph G, or max eccentricity of graph G
~~~

### Radius of gyration

Considering atoms as points in a three-dimensional space, the radius of gyration is defined as

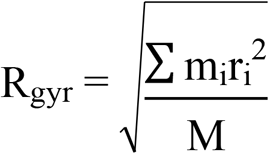

where M is the total mass of the molecule, and m_i_ is the mass of the i-th atom with distance r_i_ from the centre of mass. Radius of gyration was calculated using the *rgyr* function, which is part of the Bio3D R package (29).

### Solvent accessibility of residues

Secondary structure (total solvent-accessible area of proteins) was assigned according to the method of Kabsch and Sander (30,31). The solvent accessible area data for protein residues were taken from the work of Tien and co-workers (32).

**Table 1:**
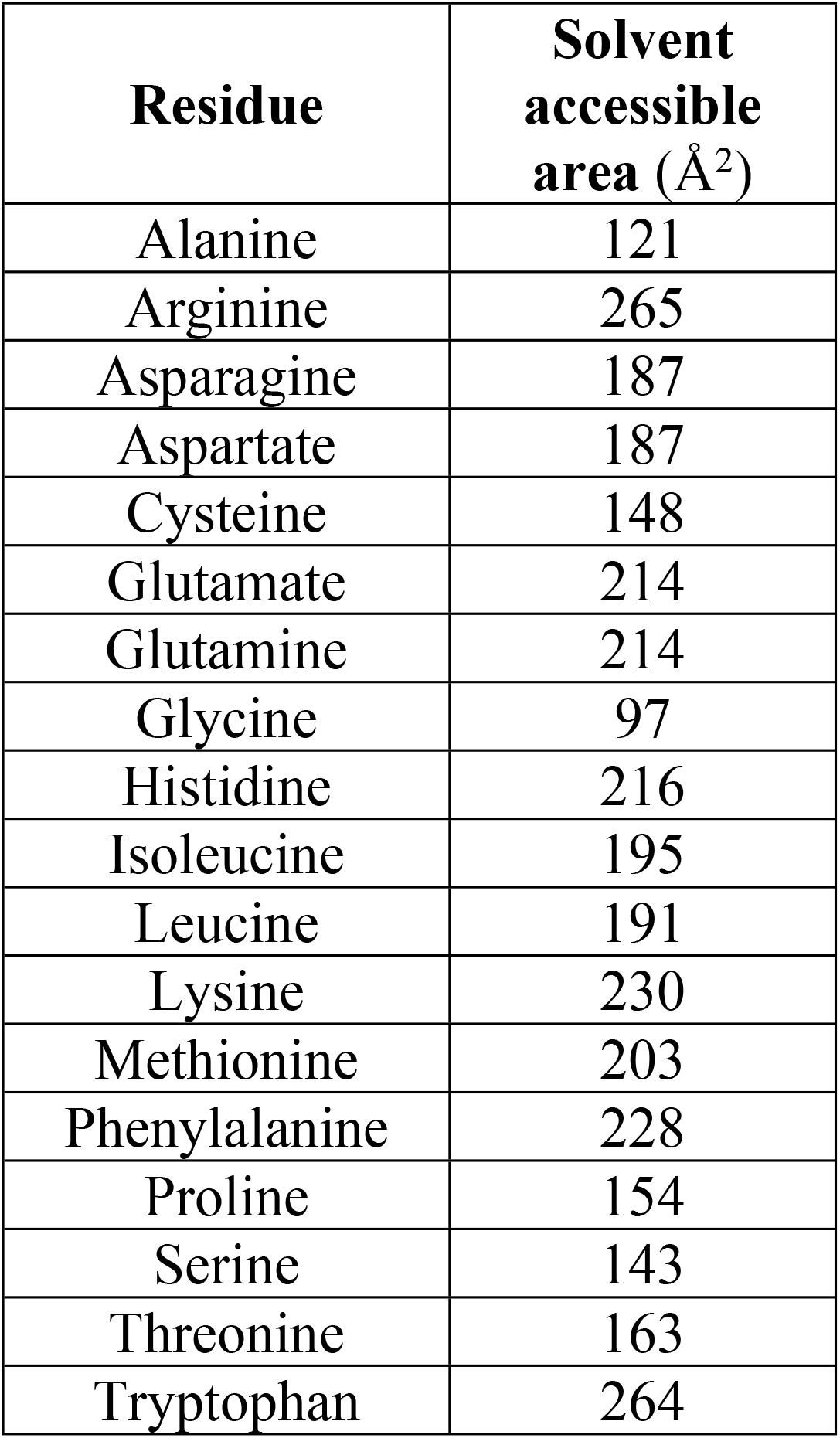
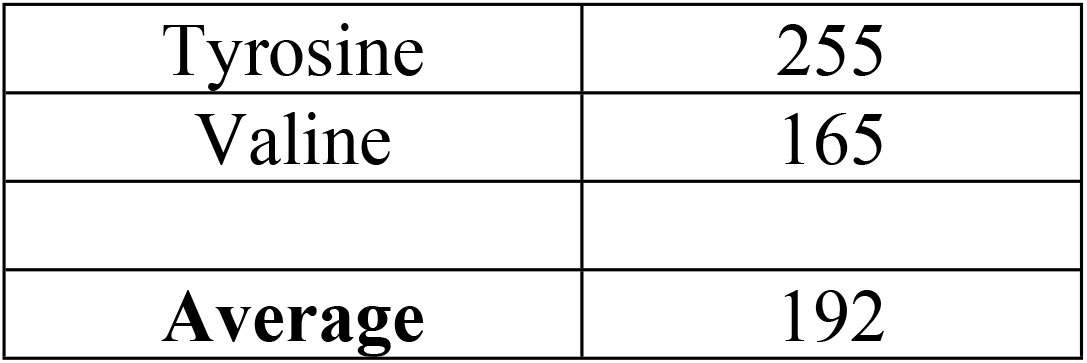
Solvent accessibility of residues.

| Residue | Solvent accessible area (Å<sup>2</sup>) |
| --- | --- |
| Alanine | 121 |
| Arginine | 265 |
| Asparagine | 187 |
| Aspartate | 187 |
| Cysteine | 148 |
| Glutamate | 214 |
| Glutamine | 214 |
| Glycine | 97 |
| Histidine | 216 |
| Isoleucine | 195 |
| Leucine | 191 |
| Lysine | 230 |
| Methionine | 203 |
| Phenylalanine | 228 |
| Proline | 154 |
| Serine | 143 |
| Threonine | 163 |
| Tryptophan | 264 |
| Tyrosine | 255 |
| Valine | 165 |
| **Average** | 192 |

## Results and Discussion

### Node degree-nearly scale-invariant

One of the fundamental global graph parameters in graph theory is the mean node degree (MND). The mean node degree shows how many edges each node has on average. In previous work, Pražnikar and co-workers (33) have shown that protein models that deviate from the expected MND by approximately two standard deviations or more are likely to be incorrect. Furthermore, the scaling exponent calculated in the mentioned study is close to zero and indicates that the mean node degree is nearly scale-invariant.

In this study, a large non-redundant database of biological units (31,571) was used, rather than crystal asymmetric units. We can see in Figure 1A that MND is not strongly dependent on protein size and that the distribution is rather narrow. Upon closer examination, however, the value determined in our study (0.038) differs slightly from the value (0.024) determined in previous study. The reason for the different scaling exponents is probably that the datasets are not the same. An analysis performed on two large but different datasets shows that MND is nearly scale-invariant, i.e., the scaling exponent is close to zero (0.024 and 0.038). Thus, MND is not strongly dependent on protein length, and it can be concluded that the number of edges in the protein graph is linearly related to the number of nodes (amino acids).

**Fig 1.**
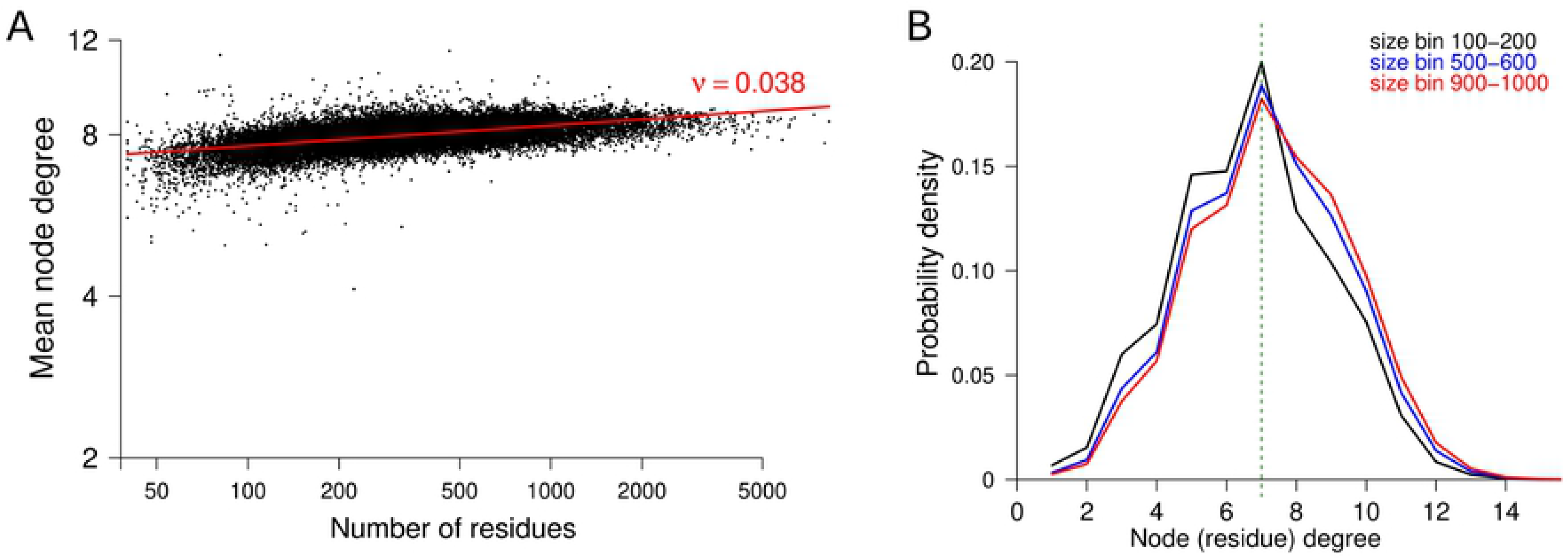
(A) Scaling exponent of mean node degree of protein graphs versus the number of residues. (B) Probability of node degree of protein graphs for three size bins. The first size bin (black line) encompasses protein structures with length between 100 and 200 residues, the second size bin (blue line) encompasses protein structures with length between 500 and 600 residues, and the third size bin encompasses structures with length between 900 and 1000 residues.

We could expect that larger proteins would have a higher average node degree because of a higher number of core residues and a relatively lower number of surface residues, which are supposed to have lower numbers of edges. Thus, to further analyse the node degree of protein nodes-residues, we calculated the node degree distribution for three different size bins. The first size bin contains all protein structures from the database, which have lengths between 100 and 200 residues, in the second size bin are proteins with lengths between 500 and 600 residues, and in the third size bin are structures with lengths between 900 and 1,000 residues. Figure 1B shows that all three distributions are very similar and that there is a peak at 7 node degrees. The comparison of all three peak values shows that the first size bin, which contains the smallest proteins, has the highest probability density value. The lowest probability density value at 7 node degree is seen for the third size bin, which contains the largest proteins among all three selected size bins. A closer look at the left (degree 2) and right (degree 14) tail of the distributions shows high similarity for all three distributions. The visual comparison shows the most significant differences on the left and right sides of the peak. The differences can be observed at values of approximately 5 to 9 node degrees. It can be seen that the first size bin has a higher probability at 3 to 6 node degrees as compared to size bins two and three. The order is somehow reversed on the right side of the displayed distribution. For node degrees 8, 9 and 10, size bin three exhibits higher values compared with size bins two and three.

This analysis shows that despite the different sizes of the proteins, they have a very similar node degree distribution, which peaks at node degree 7. A simple way to explain the presented results is that buried residues in small or large proteins form approximately the same number of links. This is a direct consequence of the fact that the amino acids are physical objects and cannot be arbitrarily close to each other. The marginal difference in node degree distributions, a slight shift to higher node degrees, explains the low positive scaling exponent (0.038), which is nearly scale-invariant.

### Protein graph eccentricity: an alternative method for analysis of radius of gyration

It is easy to ask a question: radius of gyration and radius as a graph parameter have a common name, but do they follow the same power law? To answer this question, linear regression analysis and scaling exponent were calculated for three graph parameters: radius, diameter and mean eccentricity. The radius-graph parameter is defined as minimum eccentricity, whereas the eccentricity of the graph is defined as the maximum distance between one node and all other nodes. Notice that the diameter is defined as maximum eccentricity.

Figure 2A shows the scaling exponent of a radius of gyration for 31,571 selected protein structures. The non-linear fitting function can be written as

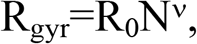

where R_gyr_ is the radius of gyration, R_0_ is the pre-factor and *v* is a scaling exponent. The pre-factor R_0_ can be obtained experimentally and used as a restrained value during non-linear fitting (34–36). Thus, when restrained fitting was performed, the pre-factor (R_0_=2 Å) was fixed. We can see that in the case of restrained fitting, the scaling exponent is 0.405, which is consistent with other studies. When fitted without restraint, the exponent is lower (0.351). Both values are within the range reported by other studies (25,27,34,37–39).

**Fig 2.**
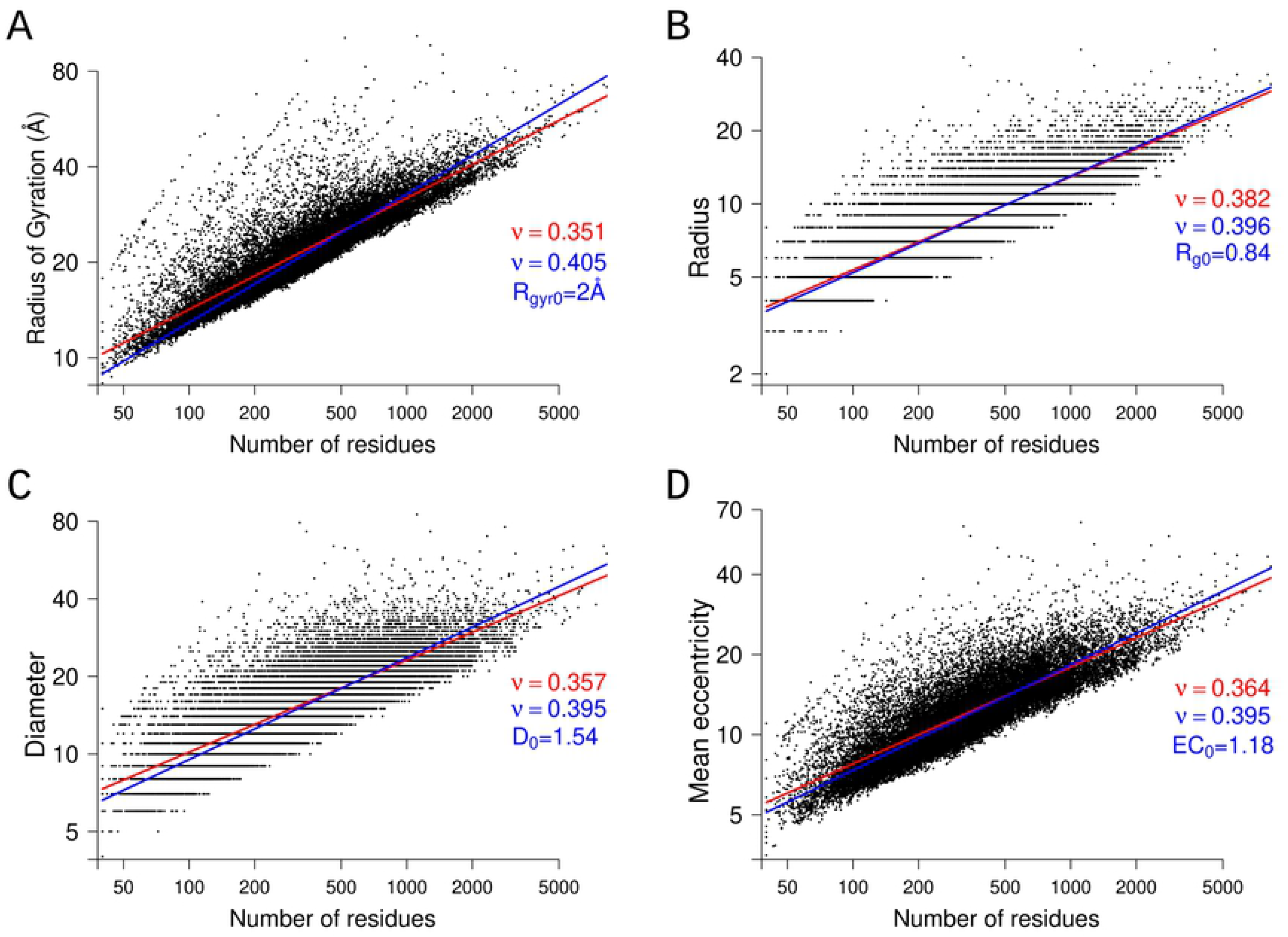
Log-log plots of the radius of gyration (A), radius (B), diameter (C), and mean eccentricity (D) versus the number of residues. The solid red line is generated by fitting without restraint; the blue line is produced by restrained fitting. The legend shows unrestrained scaling factors in red. The restrained scaling factors and corresponding pre-factors are in blue.

Figure 2B, C, and D show radius, diameter, and mean eccentricity plotted against protein length. Similar to the analysis of the radius of gyration, the power exponent was fitted with and without restraint. The pre-factor for restrained fitting was derived from linear regression analysis, as shown in Figure 3. The linear regression analysis between the radius of gyration and graph parameters reveals that R^2^ is close to 0.90 for all three cases (Fig. 3). The highest R^2^ (0.91) is observed between mean eccentricity and radius of gyration (Fig. 3C). The reason for this is probably that the values of radius and diameter are discrete, while mean eccentricity values are not discrete. For example, the radius can be 7 or 8, but cannot be a real number between 7 and 8. Mean eccentricity is just a mean value of all shortest paths to any nodes. It is seen that the distribution of mean eccentricity is smoother in comparison to the discrete values of radius and diameter on the y-axis.

**Fig 3.**
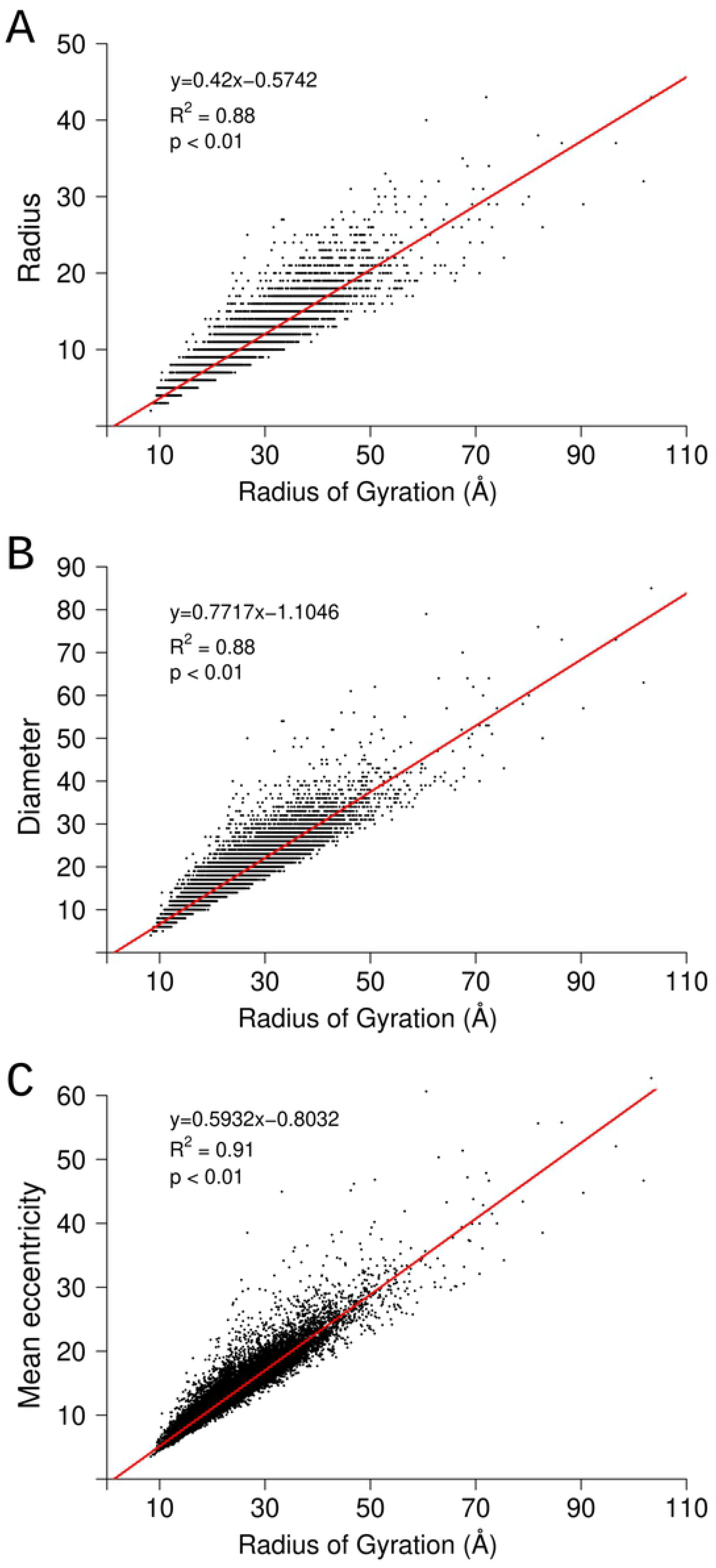
Scatter plot of graph parameters: radius (A), diameter (B), and mean eccentricity (C) versus radius of gyration. The legend shows the regression coefficient, R-squared value, and p-value.

If we use pre-factor R_0_ of a radius of gyration, which was obtained from experimental data, then we can calculate the pre-factors for radius, diameter, and mean eccentricity using the slope *k* from a regression analysis. The steepness of the linear regression model between R_gyr_ and radius was 0.42 Å^−1^, 0.77 Å^−1^ between R_gyr_ and diameter, and 0.59 Å^−1^ between R_gyr_ and mean eccentricity (see Fig. 3A, B and C). Using pre-factor R_0_ and the steepness of linear fit *k*, the pre-factors for radius, diameter and mean eccentricity can be calculated using the next expression:

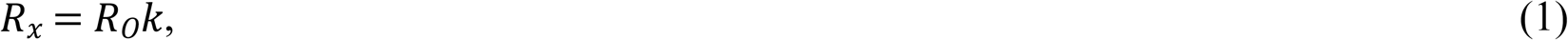

where R_x_ is a new calculated pre-factor, R_0_ is the pre-factor of radius of gyration and *k* is the steepness of the linear fit. In Figure 2B, C and D are shown calculated pre-factors for radius (2.0 Å 0.42 Å^−1^ = 0.84), diameter (2.0 Å 0.77 Å^−1^ = 1.54) and mean eccentricity (2.0 Å 0.59 Å^−1^ = 1.18). We can see that the restrained scaling exponent is higher than the non-restrained scaling exponent for all three cases. Furthermore, it is observed that restrained graph parameters all have very similar scaling exponents (~0.395) which are very close to the scaling exponent of the radius of gyration (0.405).

Thus, this study shows that the radius of gyration, which is calculated from the atomic coordinates and radius of graph follow the same scaling exponent. From this we can conclude that when analysing 3D models of macromolecules using a graph-theoretical theory approach, the eccentricity of the graph can be used to estimate the radius of gyration. Thus, graph parameter eccentricity allows us to investigate whether the 3D protein model deviates from the expected value of the radius of gyration and to obtain information about the compactness of the structure. Furthermore, when a scientist builds a model, and the model is still in an early phase, e.g., as an alanine chain, then the Cα only model, which can be represented as a graph, contains enough information to estimate the radius of gyration.

### Central and peripheral nodes

In the previous section, the relation between the radius of gyration and graph eccentricity was introduced. When exploring graph eccentricity, it is ubiquitous to examine which nodes-amino acids are central (close to every other node) and peripheral (distant from every other node). This kind of analysis has some common points with analysis of solvent-exposed residues, which is directly related to the arrangement of residues in 3D space. Buried residues constitute the core of the protein; meanwhile, residues exposed to the solvent represent the outer part of the protein in 3D space (40). The molecular mass of a protein is related with the total solvent exposed surface using the next expression:

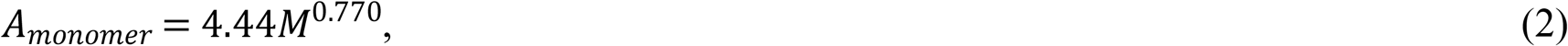

where M is the molecular mass of the protein (41). Similarly, we can introduce the relation between protein length and total solvent-exposed surface. Given that the average amino acid has a total solvent exposed area of 192Å^2^ (32), we can use this value as a restraint during data fitting. Figure 4 shows the protein length against the total exposed area. The scaling exponent of the fitted curve is 0.772, almost the same as the exponent in equation 2. This result was somehow expected because molecular mass correlates with sequence length.

**Fig 4.**
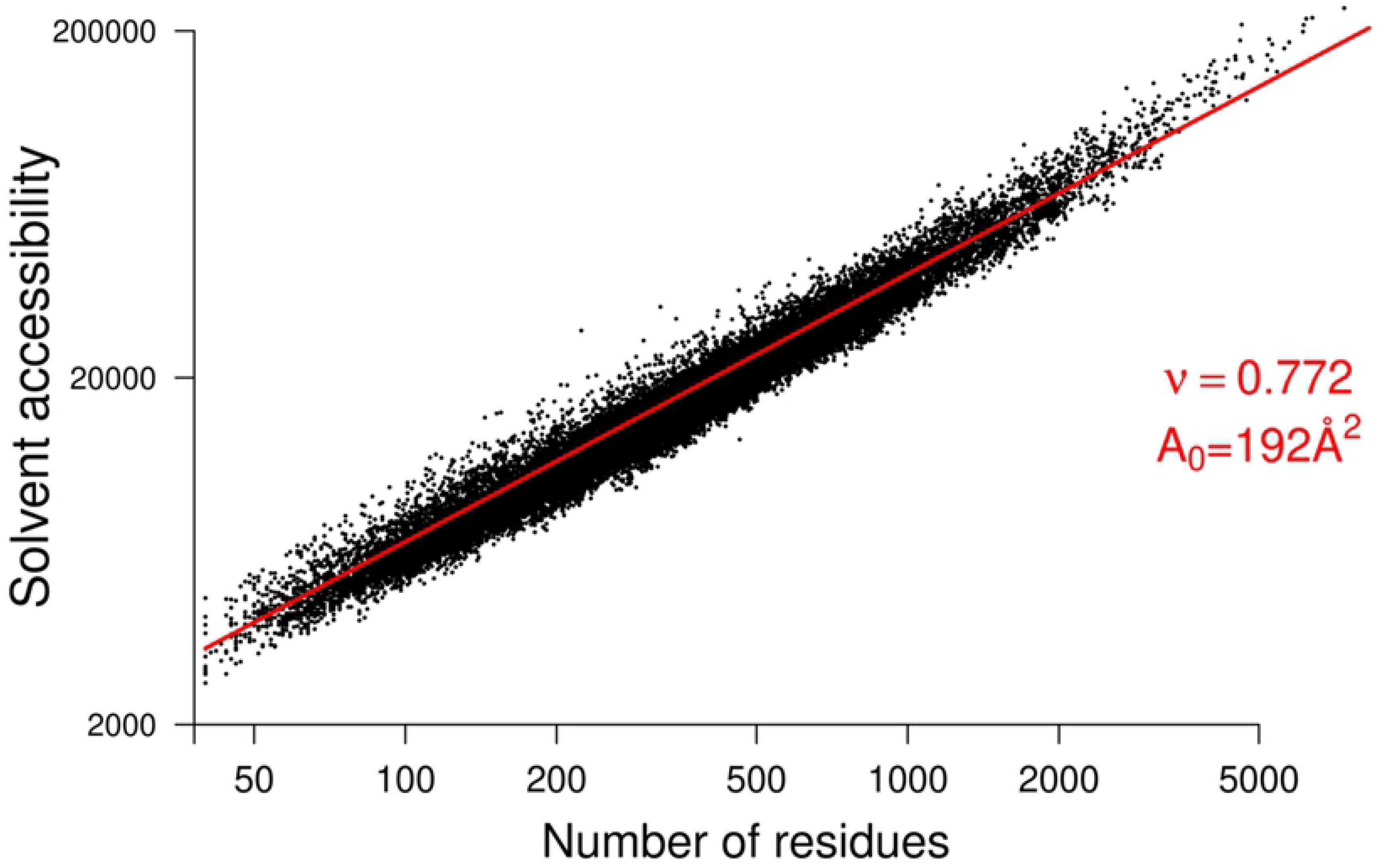
Dependence of total solvent accessible area on the number of residues in proteins. The legend shows the scaling exponent and pre-factor (A_0_), which was used during restrained fitting.

However, a closer examination of the relationship between the number of central (peripheral) nodes and protein length demonstrated that the numbers of central and peripheral residues are not related to the protein size (Figure 5A, B). To further support this finding, the comparison between the radius of gyration, mean eccentricity, and numbers of central and peripheral residues for nine different size bins was made. We can see (Figure 6) that the mean eccentricity and radius of gyration increase according to the length of the protein. Note that the scaling exponent for both mentioned parameters is approximately 0.4 (Figure 2). On the other hand, central and peripheral box plots do not show such a positive trend (Figure 6C, D). It follows that the numbers of peripheral and central residues are not related to the protein size.

**Fig 5.**
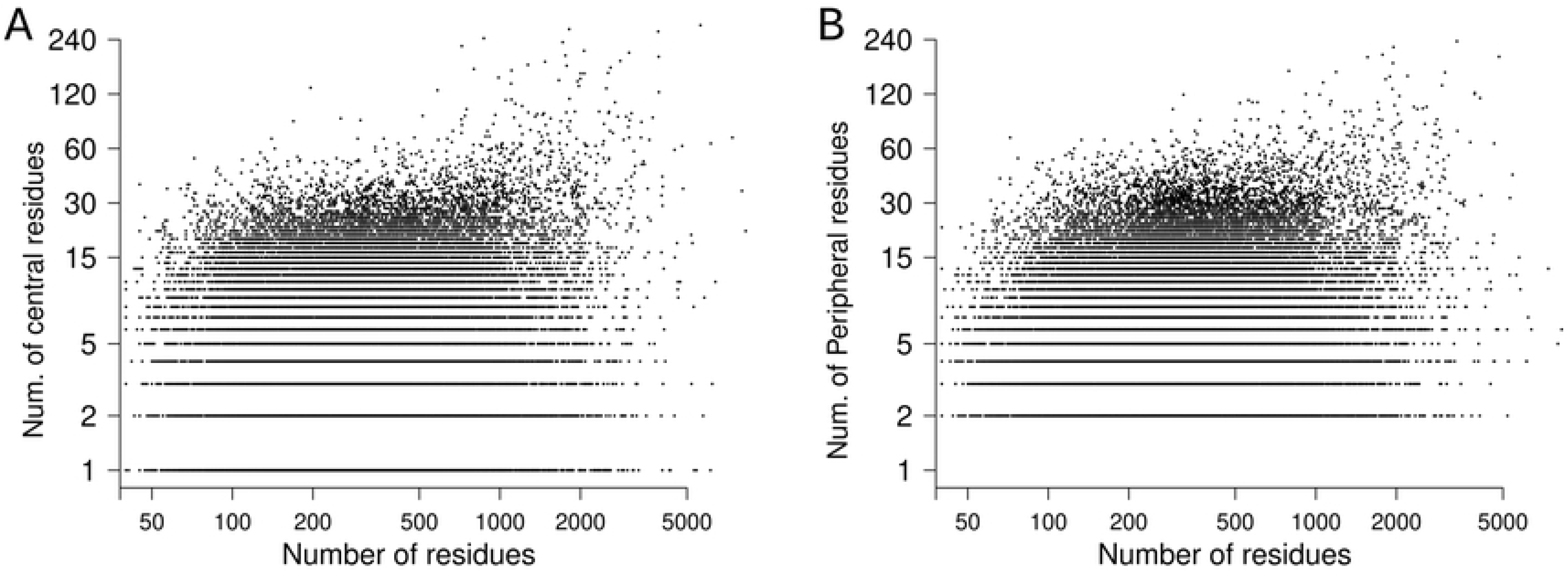
Distribution of central/peripheral residues against the size of proteins (number of residues).

**Fig 6.**
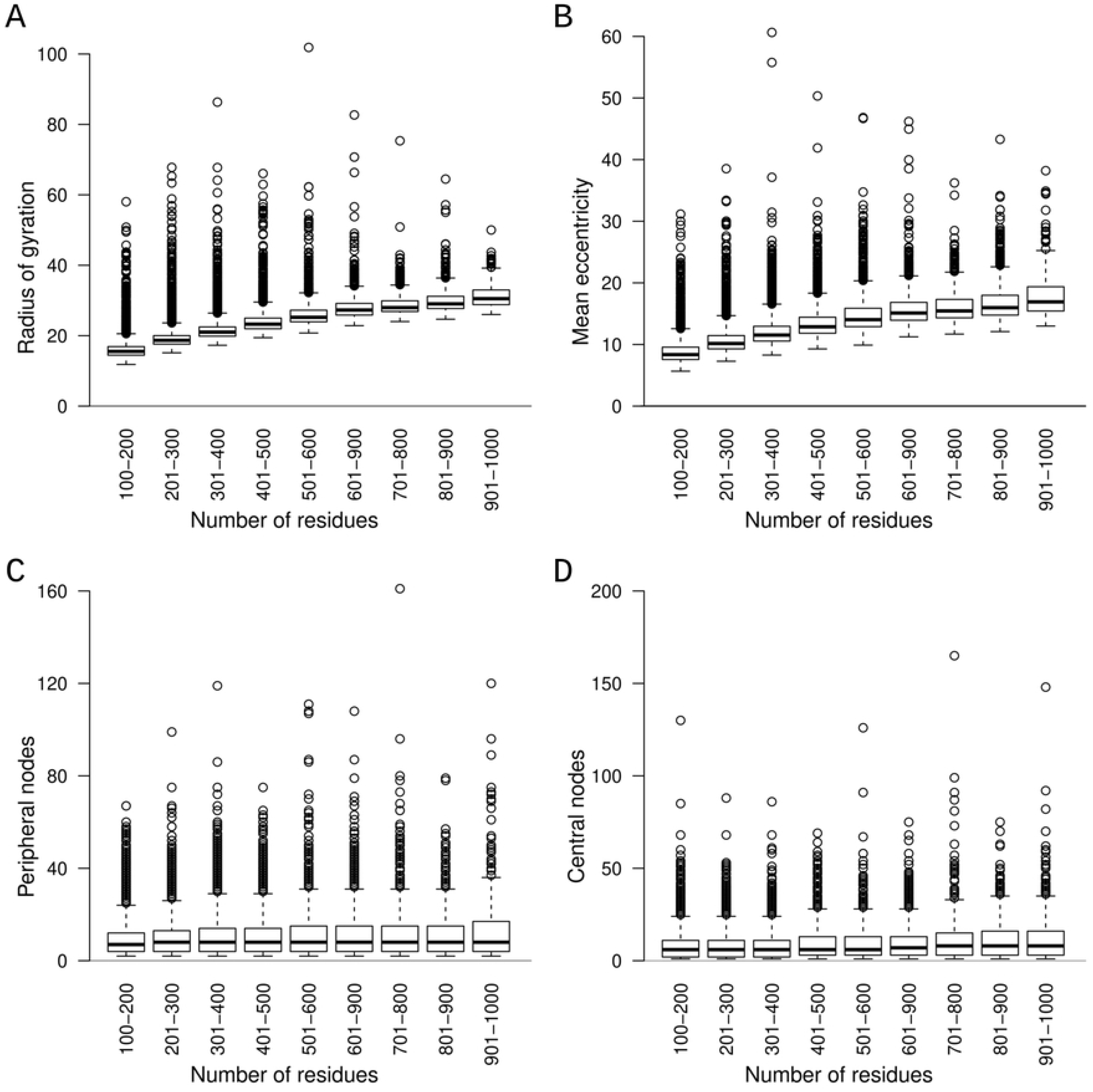
Boxplot of the radius of gyration (A), mean eccentricity (B), number of peripheral nodes (C), and number of central nodes (D) for ten different size bins.

Next, we examine the case with different central to peripheral node ratio, which is shown as a graph and ribbon representation of two protein structures (Figure 7 A, B, D, and C). The PDB id: 1pq7 structure has a low number of central nodes (4), but a high number of peripheral nodes (47): see Figures 7A and B. The situation is somehow reversed for the PDB id: 3wvj structure, which has a higher number of central nodes (34) and lower number of peripheral nodes (2): see Figure 7D, E.

**Fig 7.**
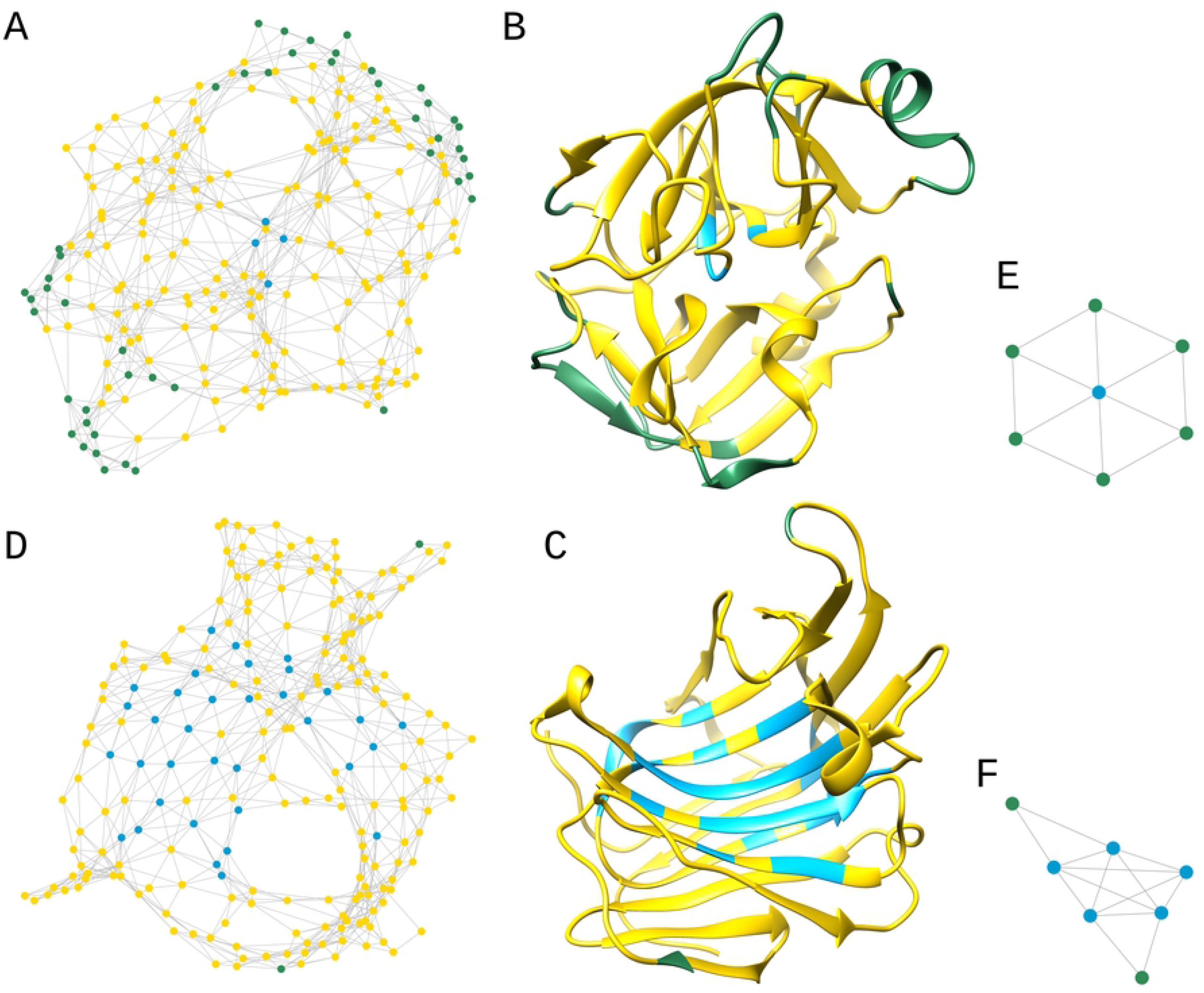
Examples of central and peripheral protein graphs. (A) Graph and (B) ribbon presentation of Biological Assembly 1 of PDB entry 1pq7. (D) Graph and (C) ribbon presentation of Biological Assembly 1 of PDB entry 3wvj. (E) Presents an almost peripheral graph, while (F) represents an almost self-centred graph.

This case shows that the numbers of central and peripheral nodes depend on the protein fold rather than on the length of the protein chain. Furthermore, we can draw parallels between *almost-peripheral* (42), *self-centred* (43), and protein graphs. Figure 7E shows the wheel, a graph that is *almost-peripheral*, containing only one central node and 6 peripheral nodes. Meanwhile, the graph shown in Figure 7F is an almost *self-centred* graph that contains 5 central nodes and 2 peripheral nodes. We could say that the graph abstracted from the PDB id: 3wvj structure is centred, while peripheral nodes-amino acids dominate the PDB id: 1pq7 structure.

Further analysis shows the frequency distribution of central and peripheral amino acids (Figure 8), which are subdivided into three groups: (i) charged, (ii) polar, and (iii) non-polar side chains. It can be seen that amino acids histidine, cysteine, methionine, and tryptophan have the lowest probability values; it follows that they are neither central nor peripheral nodes. Further, we can see that central nodes are hydrophobic amino acids (Val, Leu, Ile, Phe), which tend to be buried, while peripheral nodes are more likely hydrophilic residues (Asp, Glu, Lys) which form hydrogen bonds with solvent.

**Fig 8.**
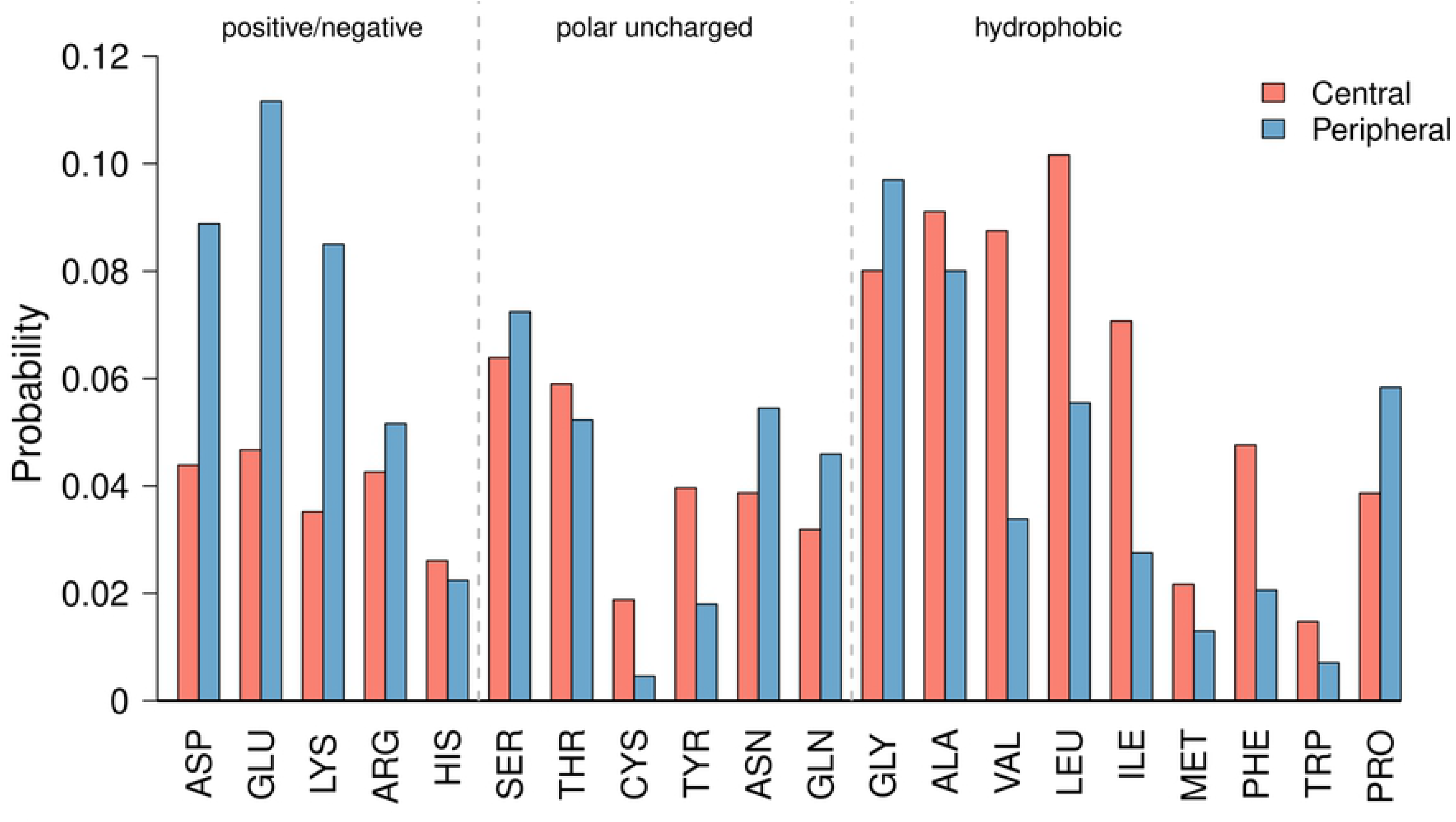
Probability density of central and peripheral nodes for different types of amino acids.

Thus, when the 3D protein structure is analysed using graph theory, the eccentricity of the graph can be very strong for evaluating the radius of gyration when studying central nodes, which tend to be hydrophobic amino acids, and peripheral nodes, which tend to be hydrophilic amino acids. It is remarkable that this graph parameter (eccentricity) allowed us to study the compactness (R_gyr_) of the protein and the arrangement of the residues (buried/not buried), making it versatile and useful for the analysis of 3D macromolecules.

## Conclusion

This study showed that the mean node degree of the protein graph is nearly scale-invariant. In other words, a small scaling exponent (0.038) indicates that the mean node degree is scale-free. Scale invariance was further supported by the analysis of node degree distribution, which showed very similar node degree distributions for proteins that differ according to size. This scale invariably offers a valuable tool for validating structures by simply counting the number of edges. Furthermore, an additional comparison between the expected node degree and node degree of a candidate could be used to explore and interpret large deviations. For example, intrinsically disordered proteins are expected to have a considerably lower mean node degree than globular proteins of the same size.

The comparison between the mean eccentricity of the graph and radius of gyration revealed a high R^2^. In other words, the mean eccentricity and radius of gyration follow the same scaling exponent (~0.4). The eccentricity of the graph, in addition to the estimation of the radius of gyration, also allows us to study the distribution of central (buried) and peripheral amino acids (non-buried). We should be aware that the mean eccentricity alone (or radius of gyration), which is used as a constraint when running molecular dynamics simulations or manually building a model, does not provide the correctness of the protein model. It is also crucial to determine how the amino acids are distributed in real space, and this can be elucidated by studying peripheral and central nodes. Thus, a single graph parameter (eccentricity) can be used to control the compactness of the macromolecule and the distribution of amino acids in 3D space, which makes it a valuable tool for analysing protein models.

## Acknowledgement

This work was supported by Structural Biology grant P1-0048 and Infrastructure programme grant I0-0035-2790, provided by the Slovenian Research Agency.

## Data availability

The data is freely available on github at https://github.com/jure-praznikar/Scaling-laws.

